# Neuronal activity drives pathway-specific depolarization of astrocyte distal processes

**DOI:** 10.1101/2021.07.03.450922

**Authors:** Moritz Armbruster, Saptarnab Naskar, Jacqueline Garcia, Mary Sommer, Elliot Kim, Yoav Adam, Philip G Haydon, Edward S Boyden, Adam E Cohen, Chris G Dulla

**Author notes:** Correspondence: correspondence should be addressed to MA or CD.

## Abstract

Astrocytes are glial cells that interact with neuronal synapses via their distal processes, where they remove glutamate and potassium (K^+^) from the extracellular space following neuronal activity. Astrocyte clearance of both glutamate and K^+^ is voltage-dependent, but astrocyte membrane potential (V_m_) has been thought to be largely invariant. As a result, these voltage-dependencies have not been considered relevant to astrocyte function. Using genetically encoded voltage indicators enabling the measurement of V_m_ at distal astrocyte processes (DAPs), we report large, rapid, focal, and pathway-specific depolarizations in DAPs during neuronal activity. These activity-dependent astrocyte depolarizations are driven by action potential-mediated presynaptic K^+^ efflux and electrogenic glutamate transporters. We find that DAP depolarization inhibits astrocyte glutamate clearance during neuronal activity, enhancing neuronal activation by glutamate. This represents a novel class of sub-cellular astrocyte membrane dynamics and a new form of astrocyte-neuron interaction.

**One Sentence Summary:** Genetically encoded voltage imaging of astrocytes shows that presynaptic neuronal activity drives focal astrocyte depolarization, contributing to activity-dependent inhibition of glutamate uptake.

## Main Text

Astrocytes are glial cells with elaborate morphological complexity, allowing close physical and functional interactions between nanometer-scale distal astrocyte processes (DAPs) and neuronal synapses. Excitatory amino acid transporters (EAATs), including GLT1 and GLAST, are concentrated in DAPs where they remove extracellular glutamate, providing spatiotemporal control of excitatory neurotransmission. The inward rectifying potassium (K^+^) channel Kir4.1 is also present in DAPs, where it mediates K^+^ buffering by allowing K^+^ influx when extracellular levels are high. Astrocytes have low membrane resistance (R_m_, ≈ 5 MΩ) and hyperpolarized membrane potentials (V_m_) (≈ -80 mV), near the K^+^ reversal potential, thanks to their expression of a cadre of K^+^ channels. This has led to the assumption that astrocytes are electrically passive and undergo only very small changes in V_m_. Whole cell recordings from astrocyte soma show that astrocyte V_m_ changes by only a few mVs during neuronal activity and returns to baseline over the course of seconds (but see (*1, 2*) for larger changes). Because of low astrocyte R_m_ and DAP morphological complexity, there is minimal spatial propagation of depolarization in astrocytes (*3*). Therefore, measurements made at the soma at best represent a heavily filtered and attenuated version of changes in V_m_ (ΔV_m_) in DAPs.

Short bursts of neuronal activity rapidly and reversibly inhibit EAAT function with synapse specificity (*4, 5*). Because EAATs are voltage-dependent (*6*), we suspected that DAP depolarization, not seen at the soma, may contribute to activity-dependent EAAT inhibition. To address this possibility, we expressed genetically encoded voltage indicators (GEVIs) in astrocytes to image astrocyte V_m_. Using this approach, we report that neuronal activity induces large, rapid, focal, and pathway-specific depolarization of astrocyte DAPs. These depolarizations are mediated predominantly by elevated extracellular K^+^ during presynaptic activity, with an additional contribution from EAAT-mediated currents. These depolarizations lead to voltage-dependent inhibition of EAATs which enhances synaptic NMDA currents. This represents a novel form of astrocyte-neuron communication mediated by rapid, spatially specific DAP depolarization and has important implication for understanding the role of astrocytes in shaping extracellular glutamate and K^+^ dynamics.

### GEVI imaging in astrocytes reveals fast, activity-dependent depolarization

We expressed the GEVIs Archon1 (*7*) or Arclight (*8*) in layer II/III mouse cortical astrocytes using AAV-mediated transduction under the control of a modified GFAP promoter(*9*). Immunohistochemical studies show robust GEVI expression co-localized with the astrocyte marker glutamine synthase (GS), but not with the neuronal marker NeuN (Fig. 1A, B). Transduced astrocytes were then morphologically reconstructed (EAAT2-tdTomato mice (*10*)). GEVI expression was seen throughout astrocyte arborizations, including in DAPs (Fig. 1C). Transduced astrocytes showed low GFAP expression and were morphological similar to un-transduced astrocytes, consistent with minimal reactive astrogliosis (Sup. Fig. 1, 2). This confirms that our approach positions GEVIs in DAPs, enabling the imaging of V_m_ throughout the astrocyte.

**Figure 1:**
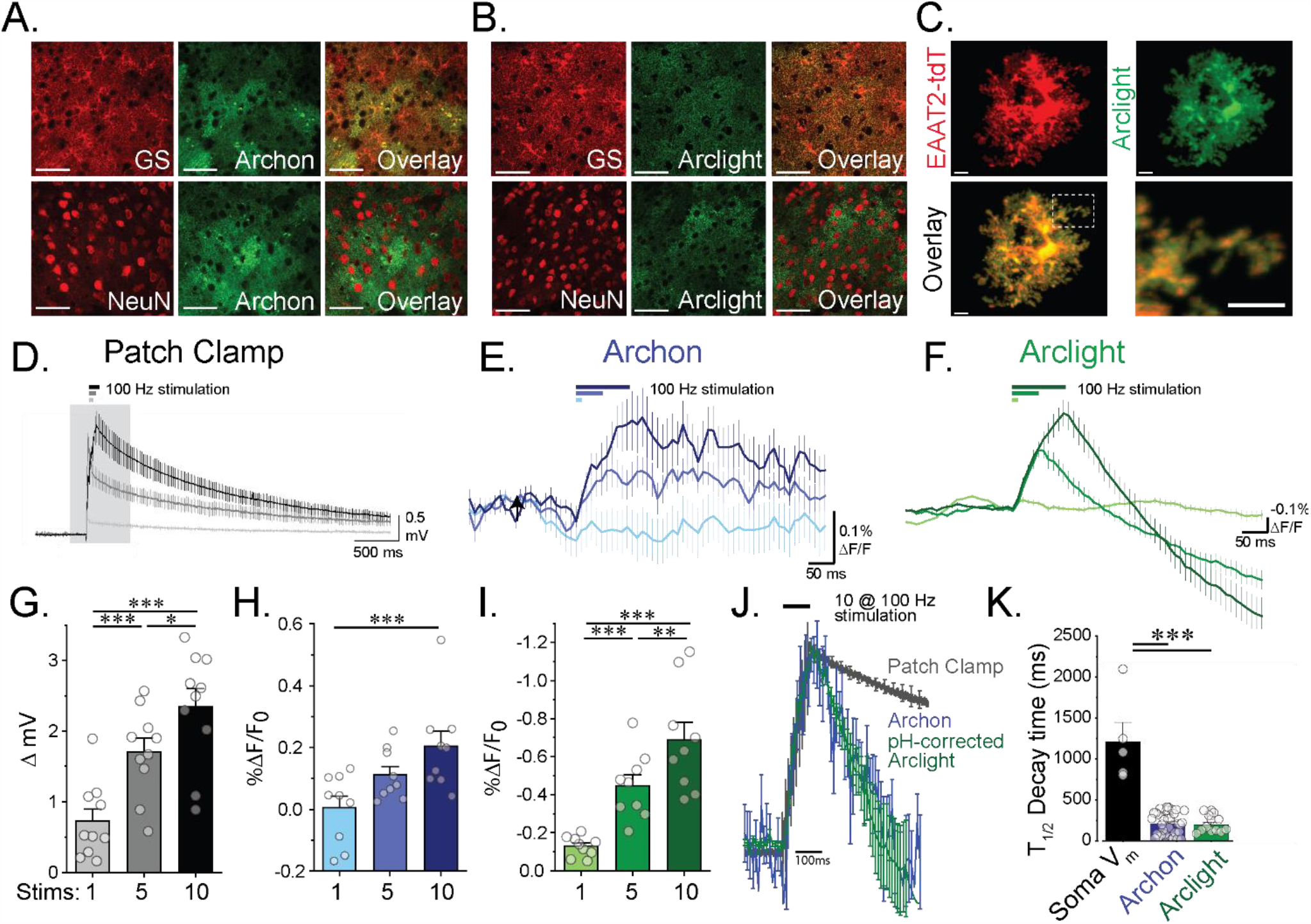
Astrocyte GEVI imaging enables measuring astrocyte DAP V_m_ changes. IHC of astrocyte expression of the GEVIs A) Archon (AAV5-GFAP-Archon) or B) Arclight (AAV5-GFAP-Arclight) show astrocyte specific expression with colocalization with the astrocyte marker glutamine synthase (GS) and lack of colocalization with the neuronal marker NeuN. Scale Bar = 50 μm C) Astrocyte reconstruction based on the astrocyte reporter line EAAT2-tdtomato, shows that Arclight GEVI labels the full astrocytic arbor. Scale bar: 5 μm, Inset 5 μm. D) Whole-cell current clamp recordings of astrocyte ΔV_m_ in response to 1, 5, or 10 Stimuli at 100Hz. Astrocyte GEVI responses to the same stimuli for E) Archon or F) Arclight. G) Whole cell recording, H) Archon, and I) Arclight show progressive depolarizations for increased stimuli number at 100Hz. 1 way repeated measures ANOVA. J) Normalized overlay of 10 Stimuli 100Hz responses of Patch Clamp, Archon, and Arclight-pH corrected decays shows significantly faster decays K) for GEVI assays compared to patch-clamp. n = 5 cells/3 mice (Patch Clamp); n = 33 slices/8 mice (Archon); n = 11 slices/3 mice (Arclight). 1 Way ANOVA.

**Figure 2:**
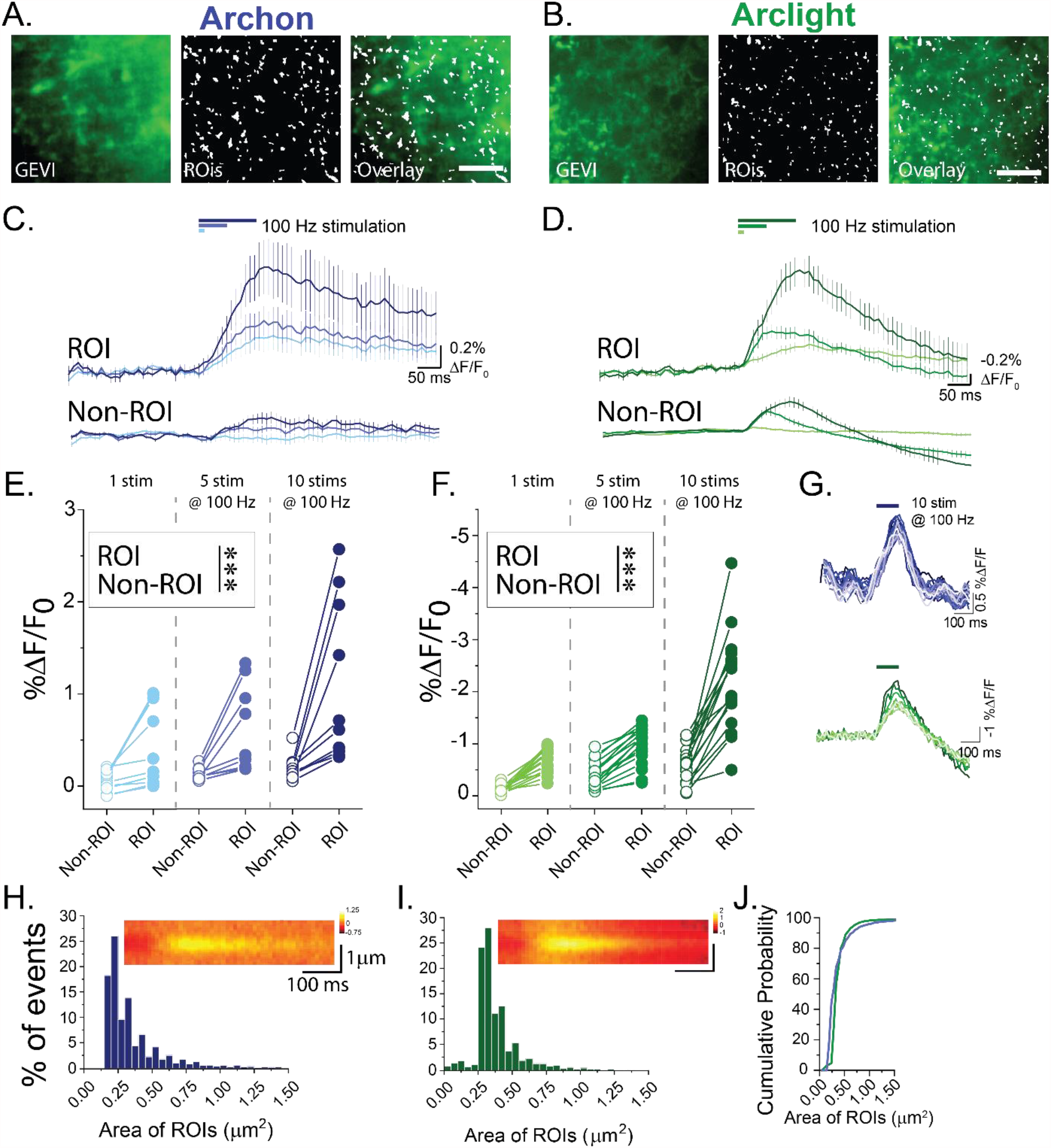
Astrocyte GEVI shows microdomain depolarizations. A) Archon GEVI colabel (GFP) and ROI maps based on 10 Stimuli 100Hz responses show small and distributed ROIs. B) Arclight (basal fluorescence) and ROI maps shows small and distributed ROIs. Scale bar = 10 µm. Average traces of ROI and Non-ROI regions from C) Archon and D) Arclight in response to 1, 5, and 10 Stimuli at 100Hz. E) Archon and F) Arclight ROIs show significantly enhanced responses (ΔF/F_0_). n = 10 slices/5 mice (Archon), n = 17 slices/6 mice (Arclight). Two way repeated ANOVA *** = p<0.001 G) Single slice examples of repeated 10 Stimuli 100Hz trains shows stable GEVI responses across multiple rounds of stimulation. Distribution of ROI sizes for H) Archon and I) Arclight shows small responsive areas and Average kymographs shows spatially restricted areas of depolarization. J) Both GEVIs show similar size distributions.

Acute coronal brain slices were then imaged on a spinning disk confocal microscope. Ascending axons were activated via electrical stimulation (1, 5, or 10 stimuli at 100 Hz) and changes in GEVI ΔF/F_0_ were quantified. Both Archon1 and Arclight showed stimulus-evoked changes in ΔF/F_0_, consistent with astrocytic depolarization (Archon1: ↑ ΔF/F_0_, Arclight: ↓ ΔF/F_0_ = Depolarization). GEVI ΔF/F_0_ amplitude increased with stimuli number (Fig. 1E, F, H, I). Membrane targeted probes predominantly report activity in astrocyte processes (*11*) as most of the astrocyte membrane is found in processes. Therefore, GEVI signal is likely heavily biased to astrocyte process V_m_. We also quantified stimulus-evoked changes in astrocyte V_m_ recorded at the soma using whole-cell electrophysiology (Fig. 1D).

We next examined the decay time of DAP depolarization monitored using GEVIs, versus those recorded electrophysiologically at the soma. Because Arclight is based on pHluorin, it undergoes pH-dependent quenching of fluorescence (*12*), interfering with quantification of ΔV_m_ kinetics. To correct this pH-effect, pHluorin quenching was imaged in separate experiments using a membrane-targeted pHluorin construct (AAV5-GFAP-Lyn-mCherry-pHluorin)(*13*). pHluorin imaging revealed pH-dependent changes in ΔF/F_0_ that were used to correct Arclight signals for changes in pH (Sup. Fig. 3). Both Archon1 and pH-corrected Arclight showed ΔV_m_ T_1/2_ decay of approximately 200 ms (Archon: T_1/2_ = 211.2 ± 24.3 ms, n = 33 slices/8 mice; Arclight: T_1/2_ = 197.3 ± 30.9 ms, n = 11 slices/3 mice), while T_1/2_ of depolarization measured with somatic whole cell recording was ≈ 5-fold slower (1206.8 ± 234.7 ms, n = 5 cells/3 mice, Fig. 1J, K).

**Figure 3:**
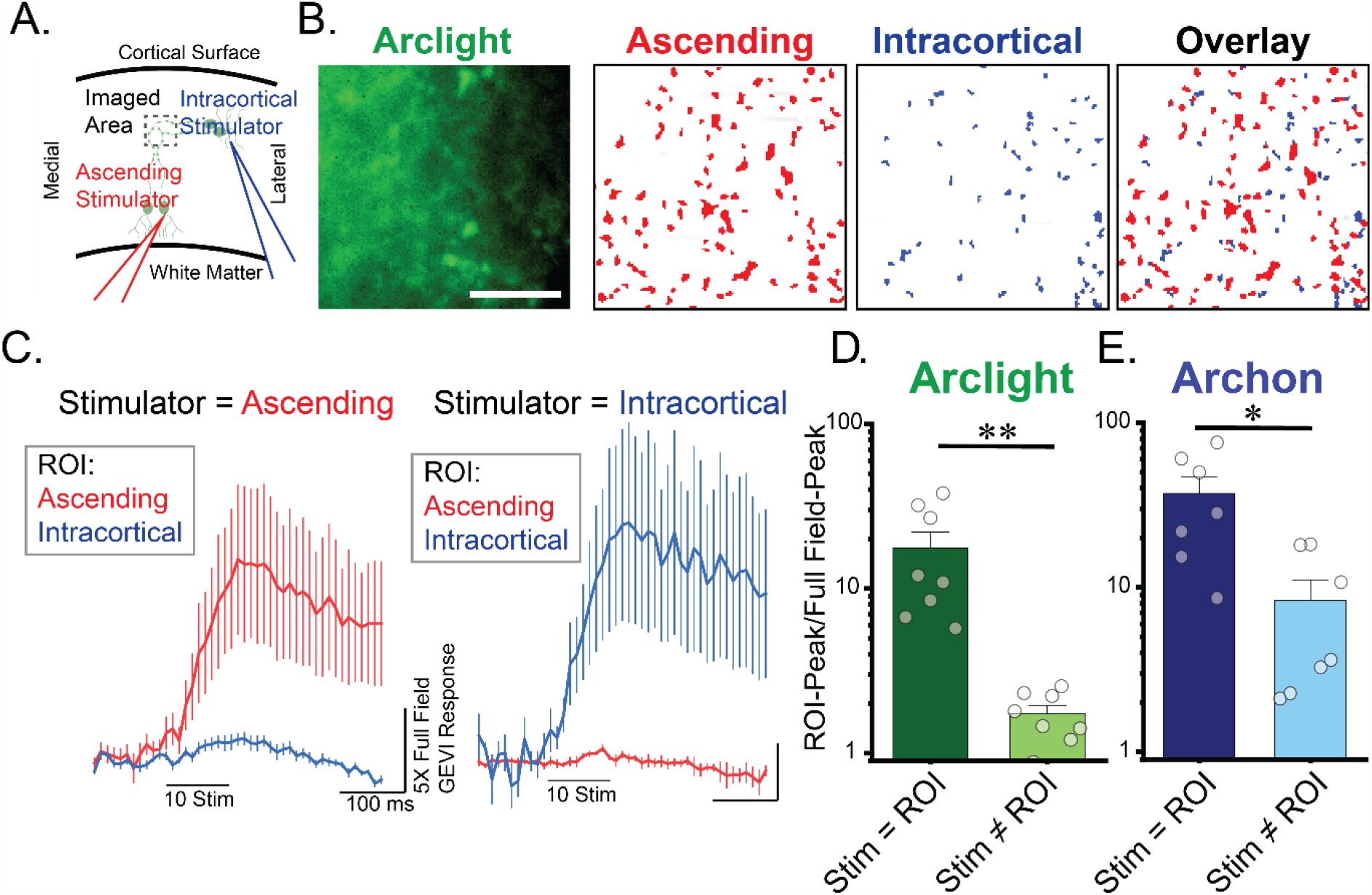
Pathway independence of astrocyte depolarization. A) To test pathway specificity we utilized an ascending stimulator (Red) and an intracortical stimulator (Blue). B) Example ROI maps of Ascending and Intracortical stimulators show little overlap. Scale bare = 10 µm. C) Average Arclight traces of matched ROI (Stim: Ascending and ROI: Ascending) or (Stim: Intracortical and ROI: Intracortical) or unmatched. D) Arclight and E) Archon matched ROI shows significant enhancement of DAP ΔF/F_0_ over unmatched suggesting pathway specificity. Log-scale. n = 8 slices/ 3 mice (Arclight); n = 7 slice/3 mice (Archon) Paired t-test * = p<0.05; ** = p<0.01.

### Activity-induced astrocyte depolarizations are focal and spatially stable

We next used principal component analysis/independent component analysis (PCA/ICA) (*14*) to identify regions of depolarization in astrocytes using both Archon1 and Arclight. This identified small “hotspots” that were used as regions of interest (ROIs) for GEVI analysis (Fig. 2A, B, Sup. Fig. 4, 5, supplemental methods). Stimulus-evoked ΔF/F_0_ in these hotspots was significantly enhanced, compared to all other regions within the imaged area (Fig. 2C-F) and were stable over repeated trials (Fig. 2G). Detected ROIs were extremely focal (Archon: 0.36 ± 0.02 µm^2^, n=10 slices/5 mice; Arclight: 0.49 ± 0.15 µm^2^, n=17 slices/6 mice), with similar size distributions for both GEVIs (Fig. 2H-J). Kymographs also confirmed spatially restricted depolarizations (Arclight: FWHM = 446±7 nm, n = 8 slices/5 mice; Archon: FWHM = 572 ± 17 nm, n = 9 slices/4 mice, Fig. 2H-I).

**Figure 4:**
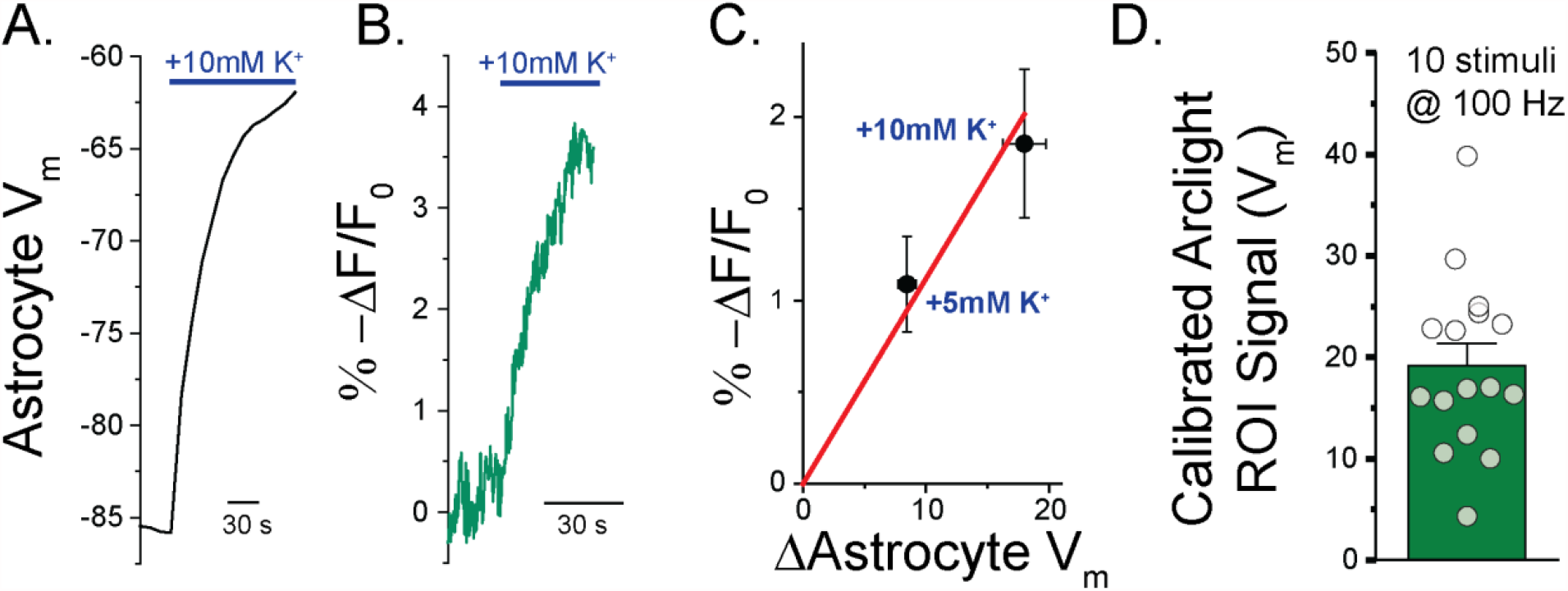
Calibrating Arclight-GEVI. Example traces of A) Astrocyte whole-cell recordings and B) Arclight GEVI show depolarizations in response to addition of high K^+^ to the aCSF (+10mM K^+^). C) Correlating whole-cell ΔV_m_ changes and Arclight ΔF/F_0_, in response to addition of +5mM and +10mM K^+^ to the aCSF solutions and fit with a linear-fixed slope trace (red). Arclight N= 7 slices/3 animals per condition, Whole-cell 8 cells/3 animals per condition. D) correcting peak ROI 10 stimuli 100Hz Arclight responses to ΔV_m_ using the calibration. N= 17 slices/6 mice.

**Figure 5:**
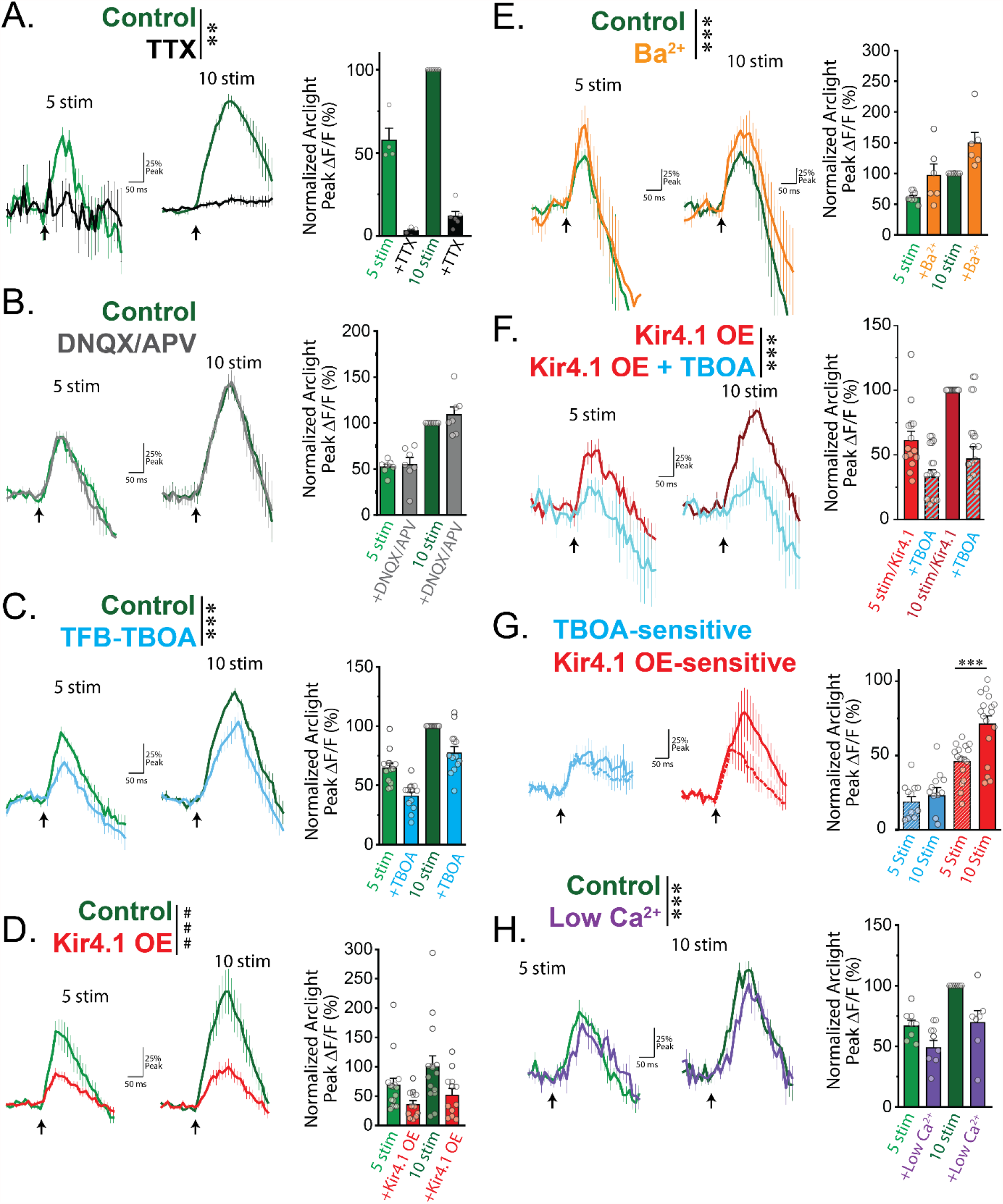
Glutamate transport and increases in [K^+^]_e_ contribute to astrocyte depolarization. Arclight GEVI imaging in response to 5 and 10 stimuli at 100Hz. A) TTX completely abolishes the GEVI response. n = 6 slices/3 mice. B) Post-synaptic glutamate receptors DNQX and APV have no effect on the peak GEVI response. n = 7 slices/3 mice. C) The EAAT inhibitor TFB-TBOA reduces the peak GEVI signal for 5 and 10 stimuli. n = 13 slices/3 mice. D) Overexpressing the astrocytic potassium channel Kir4.1, Kir4.1-OE, significantly reduces the GEVI depolarization peak. n = 15 slices/5 mice (control); n = 11 slices/4 mice (Kir4.1-OE). E) Inhibiting Kir4.1 with 200uM Ba^2+^ enhances the GEVI signal. n = 6 slices/3 mice. F) TFB-TBOA reduces the Arclight GEVI peaks of Kir4.1-OE astrocytes, showing the effects are additive. n = 13 slices/3 mice. G) Average TFB-TBOA-sensitive and Kir4.1-OE-sensitive traces. TFB-TBOA-sensitive effect does not from 5 to 10 stimuli, while the Kir4.1-OE-sensitive effects are enhanced with additional stimuli. I) Reducing extracellular Ca^2+^, reduces the peak Arclight signal. n = 7 slices/3 mice. Two Way ANOVA ### = p<0.001. Two Way Repeated Measures ANOVA ** = p<0.01, *** = p<0.001.

### Astrocyte depolarizations are pathway specific

We next performed Arclight GEVI imaging while delivering electrical stimulation to either ascending cortical axons or LII/III intracortical axons (Fig. 3A)(*4*). Hotspots were identified for each stimulation pathway using PCA/ICA. Interestingly, there was minimal spatial overlap of astrocyte depolarization hotspots evoked by ascending and intracortical axons (Fig. 3B). ROIs identified from simulation of ascending axons showed minimal responses when intracortical axons were stimulated, and vice-versa (Fig. 3C-D). Archon1 imaging replicated the pathway-specific enrichment of ΔF/F_0_ results (Fig. 3E). Together, this demonstrates that stimulus-induced astrocyte depolarization is pathway specific and could provide a mechanism to drive synapse-specific modulation of glutamate uptake and other astrocyte functions (*4*).

### Calibrating GEVI signal in DAPs

We next estimated the ΔV_m_ associated with ΔF/F_0_ using a straight-forward calibration approach. This and subsequent experiments were performed using exclusively Arclight, due to its better signal-to-noise characteristics. Both somatic V_m_, measured electrophysiologically, and GEVI fluorescence were monitored while increasing [K^+^] in the extracellular solution. This should induce a spatially uniform depolarization of astrocyte processes and soma. As predicted, increasing extracellular [K^+^] depolarized somatic V_m_ (Fig. 4A) and altered GEVI fluorescence (Fig. 4B). We performed a linear fit of the Arclight ΔF/F_0_ and somatic ΔV_m_ in response to increasing extracellular [K^+^] by 5 mM and 10 mM (Linear fit with fixed intercept R^2^ = 0.988). Using this calibration, we estimate that the ΔF/F_0_ seen in PCA/ICA-identified hotspots reflects depolarizations of 19.2 ± 2.1 mV in response to 10 stimuli at 100Hz, approximately 10-fold more than is seen at the soma (Fig. 1D). The same calibration approach suggest that non-hotspot regions depolarize by 5.1 ± 0.8 mV (n = 17 slices/6 mice).

### Presynaptic neuronal activity drives astrocyte depolarization via elevated extracellular K^+^ and EAAT activity

To determine the mechanisms that drive DAP depolarization, we probed the effects of neuronal activity, post-synaptic glutamate receptor activity, EAAT activity, and modulating K^+^ homeostasis on GEVI ΔF/F_0_. Tetrodotoxin, which blocks voltage-gated sodium channels, eliminated stimulus-evoked GEVI ΔF/F_0_, confirming that neuronal activity is required (Fig. 5A). This also eliminated the possibility that electrical stimulation acts directly on DAPs to drive their depolarization. We next assayed the role of glutamate receptor activation in DAP depolarization by blocking both AMPA and NMDA receptors (DNQX 20 µM, AP-5 50 µM, respectively). This had no effect on GEVI ΔF/F_0,_ showing that AMPA and NDMA receptor activation does not contribute to DAP depolarization in this setting (Fig. 5B). Next, we tested whether EAAT-mediated glutamate uptake, which carries an inward, depolarizing current, contributes to activity-induced DAP depolarization. Blocking EAATs with TFB-TBOA (1 µM) partially reduced GEVI ΔF/F_0_ (Fig 5C). Interestingly, the effect of EAAT blockade was similar for 5 and 10 stimuli (Fig. 5G), consistent with suppression of glutamate release during prolonged trains of neuronal activity (*4, 5*). This also suggests that other mechanisms drive the increased GEVI ΔF/F_0_ seen with increasing number of stimuli.

Because astrocyte V_m_ is highly dependent on [K^+^]_e_ and neuronal activity increases [K^+^]_e_ (*15*), we tested whether manipulating astrocyte K^+^ handling alters activity-dependent DAP depolarization. The astrocytic inwardly rectifying K^+^ channel, Kir4.1, is the primary mediator of activity-dependent astrocyte K^+^ buffering (*16*) and can be blocked with 200 µM Ba^2+^ (*17*). Viral overexpression of Kir4.1 (Kir4.1-OE) (*18-20*) (AAV5-GFAP-Kir4.1-mCherry or AAV5-GFAP-Kir4.1-EGFP, Sup. Fig. 6) significantly reduced GEVI ΔF/F_0_ for both 5 and 10 stimuli (Fig. 5D). Unlike the effects of EAAT inhibition, the effect size of Kir4.1 overexpression on GEVI ΔF/F_0_ was significantly larger for 10 stimuli, as compared to 5 stimuli (Fig. 5G). Conversely, inhibiting Kir4.1 with Ba^2+^ caused a small but significant increase in GEVI ΔF/F_0_ (Fig 5E), suggesting that Kir4.1-mediated K^+^ influx helps to minimize activity dependent DAP depolarization. Interestingly, Ba^2+^ blockade of Kir4.1 reduces stimulus-evoked depolarization of the soma, suggesting that Kir4.1 activity has unique effects on DAP and somatic V_m_ (*16, 21*) (Fig. Supp. 7). We next tested whether the effects of Kir4.1-OE and TFB-TBOA were additive. TFB-TBOA reduced the activity-dependent GEVI ΔF/F_0_ even when Kir4.1 was overexpressed, confirming that EAAT activity and changes in [K^+^]_e_ represent distinct mechanisms contributing to DAP depolarization. This suggests that activity-dependent accumulation of extracellular K^+^ is the primary driver of stimulus-dependent DAP depolarization, while EAAT activity plays a secondary role.

**Figure 6:**
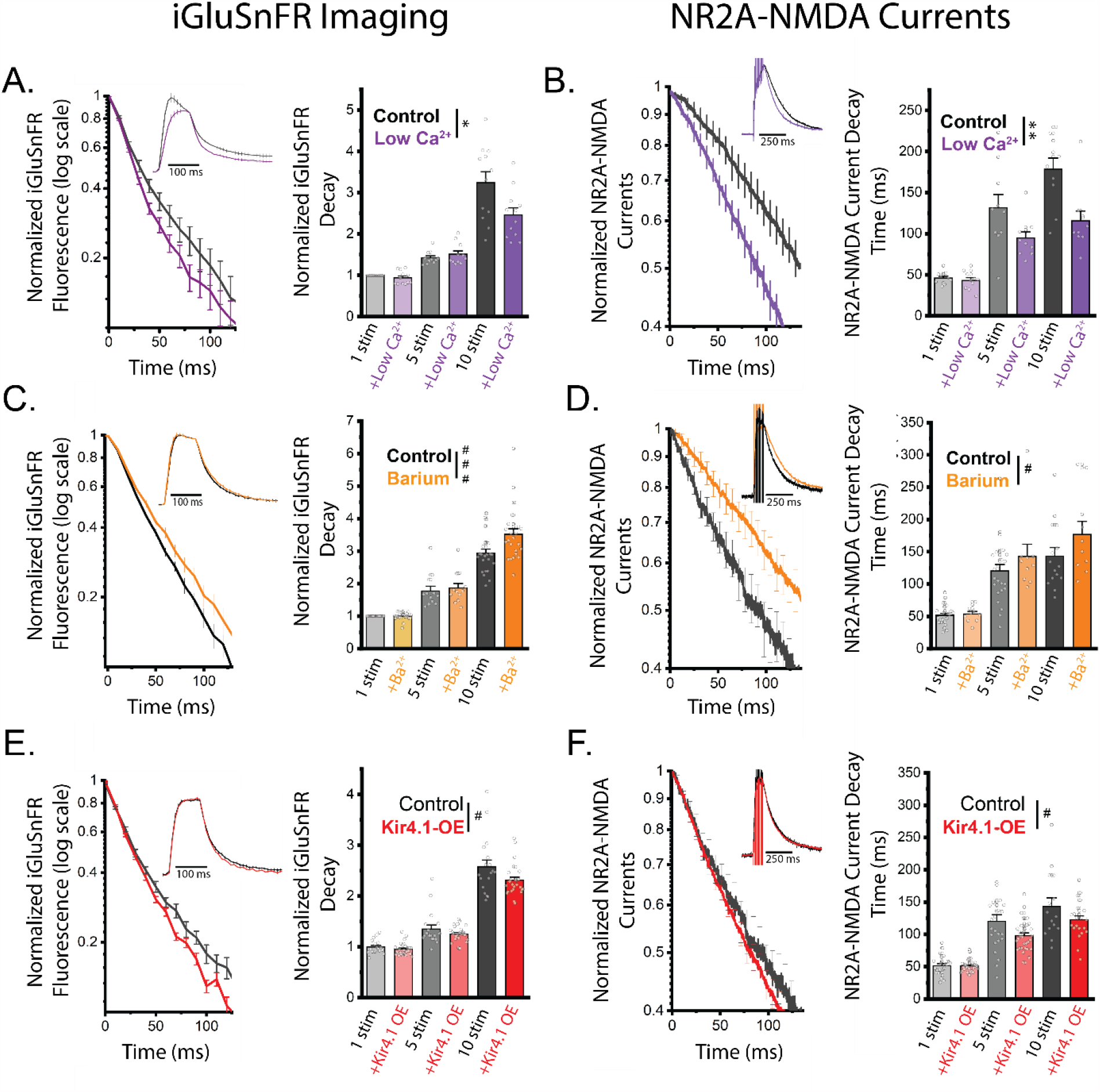
Astrocyte depolarization contributes to activity dependent glutamate clearance slowing. Glutamate clearance slowing, as assayed by single exponential fits to iGluSnFr imaging and NR2A-specific NMDA current decays in response to 1, 5 and 10 stimuli at 100Hz. A) Low Ca^2+^ accelerates average iGluSnFr decay following 10 stimuli at 100Hz. Full trace inset. n = 12 slices/5 mice B) Average 10 stimuli 100Hz NR2A-NMDA current decay shows Low Ca^2+^ aCSF speeds NR2A-NMDA current decay. Full trace inset. n = 10 cells/3 mice C) Inhibiting Kir4.1 with 200 µM Ba^2+^ slows iGluSnFr decay. n = 24 slices/6 mice D) Inhibiting Kir4.1 with 200 µM Ba^2+^ prolongs NR2A-NMDA currents. n = 15 cells/3 mice (Control); 10 cells/4 mice (Ba^2+^). E) Kir4.1-Ovexpression accelerates iGluSnFr decay. n = 18 slices/3 mice (control) n = 27 slices/4 mice (Kir4.1-OE). F) Kir4.1-Overexpression speeds NR2A-NMDA current decay. n = 33 cells/4 mice, control same as D. Two Way ANOVA # = p<0.05, ### = p<0.001. Two Way Repeated Measures ANOVA * = p<0.05, ** = p<0.01.

Finally, we asked whether reducing extracellular Ca^2+^ (which can affect presynaptic function, membrane charge screening, and Ca^2+^ signaling) altered activity-dependent changes in GEVI ΔF/F_0._ Reducing extracellular [Ca^2+^] from 2 mM to 1 mM had a strong effect on GEVI ΔF/F_0_ rise-time, due to a delayed onset of depolarization (Sigmoidal T_1/2_ rise time for 10 Stimuli 100Hz: 61.3 ± 3.6 ms Control; 85.1 ± 5.9 ms Low Ca^2+^; Paired t-test p<0.002), and caused a small but significant decrease in DAP ΔV_m_ peak (Fig 5H). While lowering Ca^2+^ decreases glutamate release, glutamate receptor activation does not drive DAP depolarization (Fig. 5B). Together, this suggests that activity dependent changes in DAPs can be altered by lowering extracellular Ca^2+^, potentially via altering astrocyte membrane charge screening(*22*) or calcium signaling.

### DAP V_m_ modulates glutamate clearance and post-synaptic NMDA receptor activation

The results we report suggests voltage-dependent modulation of astrocyte function may occur in DAPs. Astrocyte glutamate clearance by EAATs is rapid (<10 ms)(*23*), steeply inhibited by depolarization in expression systems (*6*), and slowed by neuronal activity in cortical brain slices (*4, 5*). We therefore hypothesized that EAAT function may be modulated in astrocytes by activity-induced DAP depolarization. To test this hypothesis, we used approaches that alter activity-induced DAP GEVI ΔF/F_0_ (Fig. 5) and asked whether these manipulations affect the stimulus-dependent slowing of EAAT function and glutamate clearance. Using iGluSnFr glutamate imaging (*4, 24*) and NR2A-specific NMDA currents (*4*), we tested the effects of Low Ca^2+^, Ba^2+^, and Kir4.1-OE on activity-dependent inhibition of glutamate clearance. Kir4.1-OE and Low Ca^2+^, which reduce DAP GEVI ΔF/F_0_, both reduced the slowing of glutamate clearance associated with trains of neuronal activity. The slowing of glutamate clearance is independent of the amount of glutamate released (*4*), suggesting that the effects of low Ca^2+^ are not mediated by reduced presynaptic glutamate release. Recording astrocyte glutamate transporter currents (GTCs) confirmed that lowering Ca^2+^ reduced activity-dependent inhibition of EAAT function (as assayed by GTC decay times, Sup. Fig. 8). Conversely, blocking Kir4.1 with Ba^2+^, which augments DAP GEVI ΔF/F_0_, enhanced the activity-dependent slowing of glutamate clearance. These experiments show that activity-dependent astrocyte depolarization inhibits EAAT function.

## Discussion

Using GEVI imaging, we show that DAPs undergo highly focal depolarizations (ΔV_m_ ≈ 20 mV) during brief bouts of neuronal activity. These depolarizations occur with pathway specificity, are driven by a combination of presynaptic K^+^ release and EAAT activity, and impact activity-induced slowing of glutamate uptake. These results challenge the view that astrocytes have largely invariant membrane potential and show that local astrocyte depolarizations have functional effects on EAAT activity. Our data supports that DAP V_m_ changes occur at least in part at peri-synaptic DAPs; depolarization is driven by both presynaptic neuronal activity and glutamate transport, and DAP ΔV_m_ shapes extracellular glutamate dynamics. Electron microscopy studies show that DAPs are ≈ 100 nm at sites of neuronal interaction, below the diffraction limitation of fluorescence microscopy. This limits our ability to resolve the fine spatial structure of DAP depolarization beyond our hotspot analysis. There likely exists spatial gradients in DAP V_m_ which we are not capable of quantifying due to technical constraints. Our results, therefore, may overestimate the spatial extent of DAP depolarization (FWHM ≈ 500 nm) and underestimate their peak amplitude. These caveats aside, our findings show that DAPs undergo large, rapid, and focal voltage changes during neuronal activity.

Elevation of [K^+^]_e_ appears to be a significant driver of activity-induced DAP depolarization. Astrocyte V_m_ is largely set by [K^+^]_e_, but the spatial nature of [K^+^]_e_ dynamics are not well understood. Our findings predict that neuronal activity induces highly focal elevations in [K^+^]_e_ ≈ 10 mM. This is significantly larger than generally reported during brief bursts of neuronal activity, but the small spatial scale of the predicted elevations in [K^+^]_e_ would be difficult to resolve using previous approaches like K^+^-selective electrodes. During prolonged activity (>30 s) or pathological states, like seizures, [K^+^]_e_ can increase to ≈ 10 mM, making our reported measurement physiologically possible (*15, 25*). Changes in [K^+^]_e_ and DAP V_m_ of this magnitude would have significant implications on astrocytic K^+^ buffering and may explain why Kir4.1 activity mediates unique effects on DAP and somatic V_m_. Kir4.1 takes up K^+^ when extracellular levels are elevated, thereby contributing to astrocyte hyperpolarization, but in doing so carries a counterintuitive depolarizing current. Our data suggests that DAP ΔV_m_ is primarily depolarized by elevated levels of K^+^_e_, which Kir4.1 acts to restore, thereby facilitating hyperpolarization. At the soma, however, Kir4.1 appears to contribute to astrocyte V_m_ by carrying an inward depolarizing current (*16, 21*) (Supp. Fig. 7). Therefore, the extracellular diffusion of K^+^ away from hotspots to nearby neighboring areas suggests K^+^-buffering may by highly local, with K^+^ ions moving small distances from the areas of depolarization to nearby areas that remain hyperpolarized. The prolonged depolarization seen at the soma is likely reflective of the integrated, slow K^+^ influx throughout the astrocyte and at the soma itself. Future development of K^+^_e_ imaging approaches and computational modeling of astrocytes(*26*) will continue to improve our understanding of these complexities.

Astrocytic glutamate uptake by EAATs is voltage-dependent and inhibited by neuronal activity. Our data supports that DAP V_m_ shapes EAAT function during neuronal activity. Modulating DAP V_m_, by altering [K^+^]_e_ handing, can bidirectionally shape extracellular glutamate dynamics and NMDA receptor decay times. Altering DAP V_m_ has a significant but relatively small effects on EAAT function, however, suggesting that either our manipulations do not sufficiently control DAP V_m_ or that other changes (EAAT trafficking, cell swelling, diffusion, pH changes) play a larger role in activity-dependent inhibition of EAAT function. Interestingly, neither activity-dependent EAAT inhibition nor DAP depolarization relies on glutamate receptor activation. This is surprising as post-synaptic NMDA receptor activation is thought to mediate K^+^ efflux which should depolarize astrocytes (*27*). This could represent ultrastructural differences in K^+^ handling at DAPs and soma or brain region specific astrocyte-neuron interactions. Our findings could also have important implications on astrocyte-neuron interactions and on the role astrocyte depolarization may play in activity-dependent synaptic plasticity(*28*).

How DAP depolarization affects other astrocytic functions remains to be seen, but astrocytes have complex intracellular Ca^2+^ signaling and express voltage-dependent ion channels (*29-32*), receptors (*33*), and transporters(*34*) that may be functionally modulated by V_m_. Additionally, experimental manipulations such as channelrhodopsin can increase [K^+^]_e_ (*35*) which may affect astrocyte V_m_. Finally, astrocytes are highly dynamic, responding to injury, inflammation, and more. If DAP properties are altered during reactive astrocytosis, the coupling between neuronal activity and astrocyte function may be altered, especially during behaviorally relevant bursts of neuronal activity in vivo. Together, this study shows that astrocytes experience rapid, focal, and functionally relevant depolarizations during neuronal activity.

## References and Notes

## Acknowledgements

We thank members of the Dulla, Haydon, and Rios labs, Dr. Joseph Raimondo, and Dr. Jeffery Diamond for helpful comments on the manuscript. We thank Dr. Yongjie Yang for EAAT2-tdTomato mice. We thank Dr. Loren Looger, Dr. Vincent Pieribone, Dr. Baljit Khakh, and Dr. Sergio Grinstein for making plasmids and constructs available.

## Authors Contributions

**Moritz Armbruster:** Conceptualization, Methodology, Investigation, Formal analysis, Data curation, Visualization, Writing -Original Draft. **Saptarnab Naskar:** Investigation, Formal Analysis, Writing – Review & Editing. **Jacqueline Garcia:** Investigation. **Mary Sommer:** Investigation. **Elliot Kim:** Investigation. **Yoav Adam:** Methodology, Investigation. **Phil Haydon:** Resources, Methodology. **Ed Boyden:** Resources, Methodology. **Adam Cohen:** Resources, Methodology. **Chris G. Dulla:** Conceptualization, Formal analysis, Visualization, Supervision, Funding acquisition, Project administration, Resources, Writing -Original Draft

