## Supplemental Data and Methods for "Neuronal activity drives pathway-specific depolarization of astrocyte distal processes"

**Supplemental Figure 1**

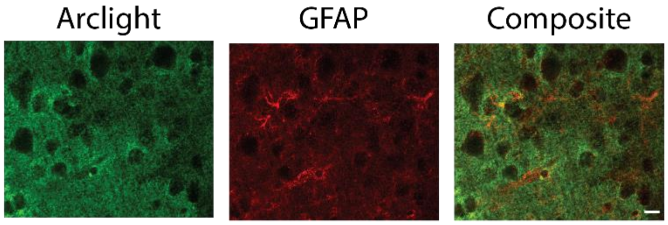


**Supplemental Figure 1: GEVI expression does not induce reactive astrocytosis.** Representative confocal IHC images stained for Arclight (green) and GFAP (red) shows the lack of reactivity in the infected astrocytes. Scalebar = 10 µm.

**Supplemental Figure 2**

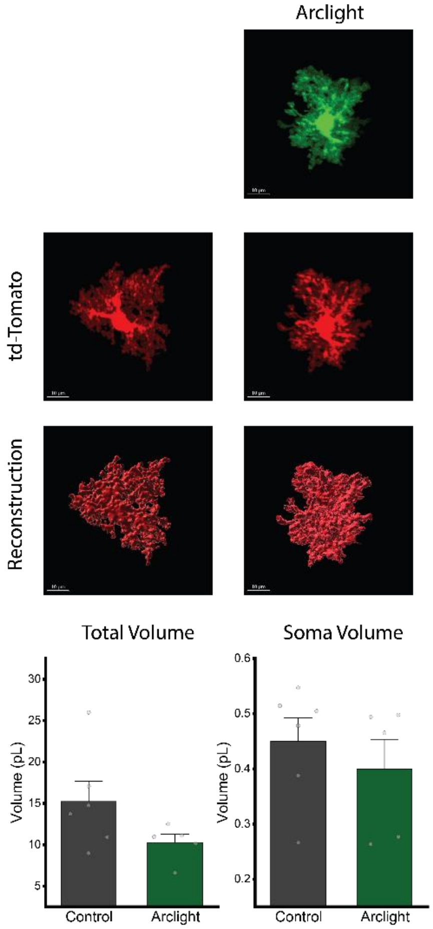


**Supplemental Figure 2: GEVI expression does not change astrocyte morphology.** Example confocal sections and reconstruction of the astrocyte reporter EAAT2-tdTomato mice either uninfected controls (left column) or Arclight infected (right column). No significant changes were observed in total astrocyte volume or soma volume between infected and uninfected astrocytes. N= 6 control, 5 Arclight astrocytes. Two-sample t-test.

**Supplemental Figure 3**

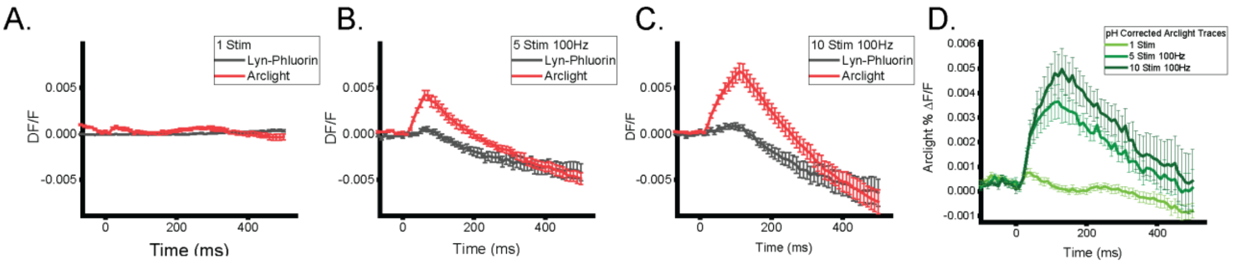


**Supplemental Figure 3: Correcting pH transients.** Average pH transients (Lyn-pHluorin) and Arclight GEVI responses to A) 1 Stim, B) 5 Stimuli 100Hz, C) 10 Stimuli 100Hz. D) The GEVI decays are corrected for the pH changes using the difference in the Arclight and pHluorin traces. n = 9 Slices/ 3 mice (pHluorin). n = 17 slices/6 mice (Arclight).

**Supplemental Figure 4**

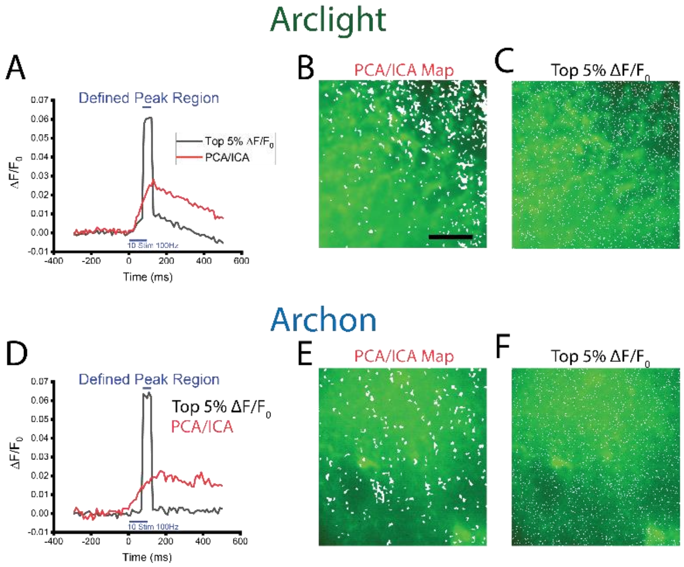


**Supplemental Figure 4: PCA/ICA ROIs are distinct from noise.** Representative examples of A) Arclight and D) Archon showing the response of PCA/ICA isolated ROIs (B, red) compared with taking the top 5% of $\Delta F/F_0$ pixels (C, black). The top 5% $\Delta F/F_0$ is defined by the peak region chosen, and shows artificial (single frame) kinetics, compared to the PCA/ICA identified ROIs. Similarly, the top 5% $\Delta F/F_0$ shows random spatial distribution (C, F), compared to the PCA/ICA maps (B, E). Scale bar = 10µm.

**Supplemental Figure 5**

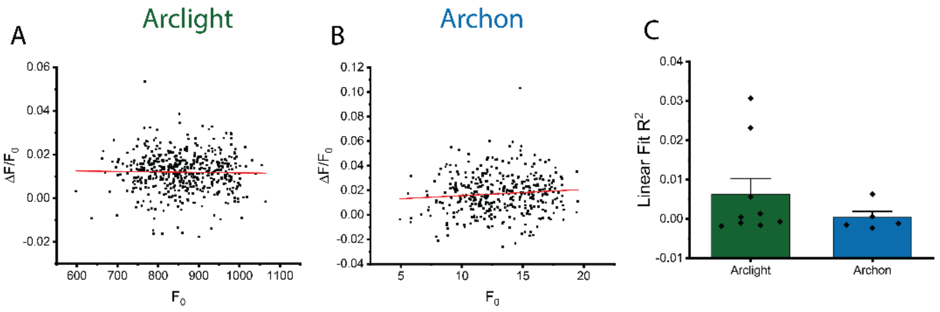


**Supplemental Figure 5: PCA/ICA ROIs are not driven by low $F_0$.** Representative examples of A) Arclight and B) Archon slices showing the distributions of individual ROIs identified by PCA/ICA analysis plotting $\Delta F/F_0$ versus $F_0$. Linear fit (red) shows no correlation between resting $F_0$ and $\Delta F/F_0$ response, suggesting that ROI identification is not biased based on the resting fluorescence. C) Average linear fit $R^2$ for Arclight and Archon are not significantly different from 0, one-sample T-test.

**Supplemental Figure 6**

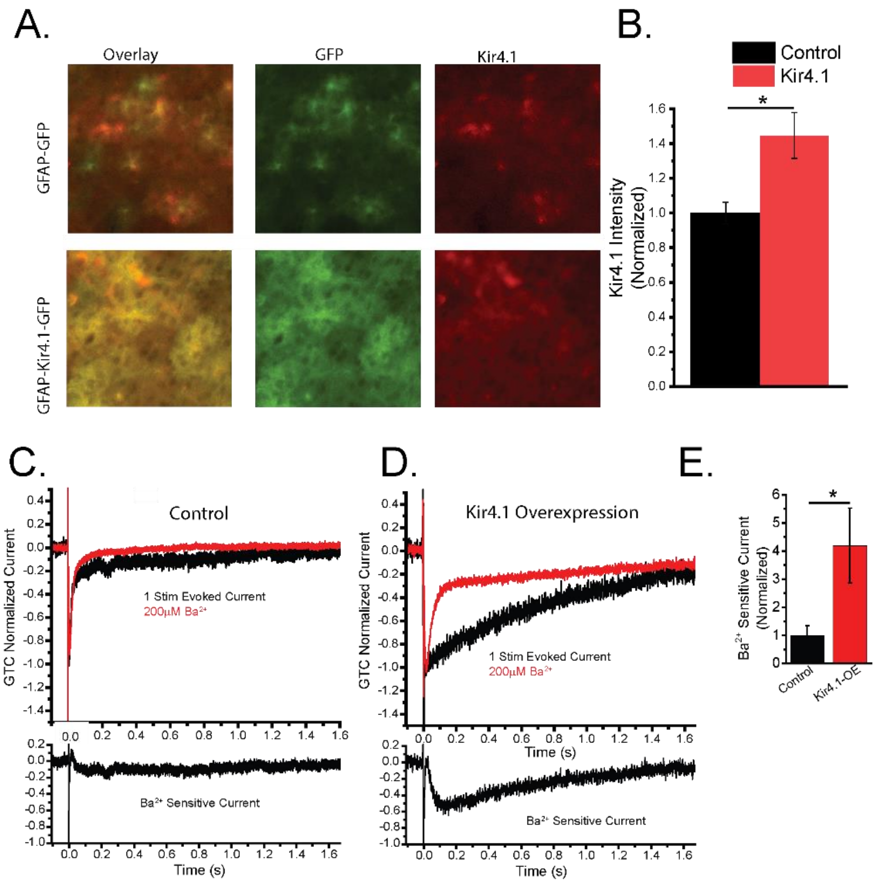


**Supplemental Figure 6: Kir4.1 Overexpression.** Epifluorescence imaging of immunofluorescence staining of Kir4.1 in Kir4.1 overexpression (Kir4.1-OE), (AAV5-GFAP-Kir4.1-EGFP) or control (AAV5-GFAP-GFP) infected cortex. Kir4.1 staining shows significant enhanced Kir4.1 staining. Scale bar = 10 µm. Two-sample t-test, n=3, 4 mice, * = $p < 0.05$.

**Supplemental Figure 7**

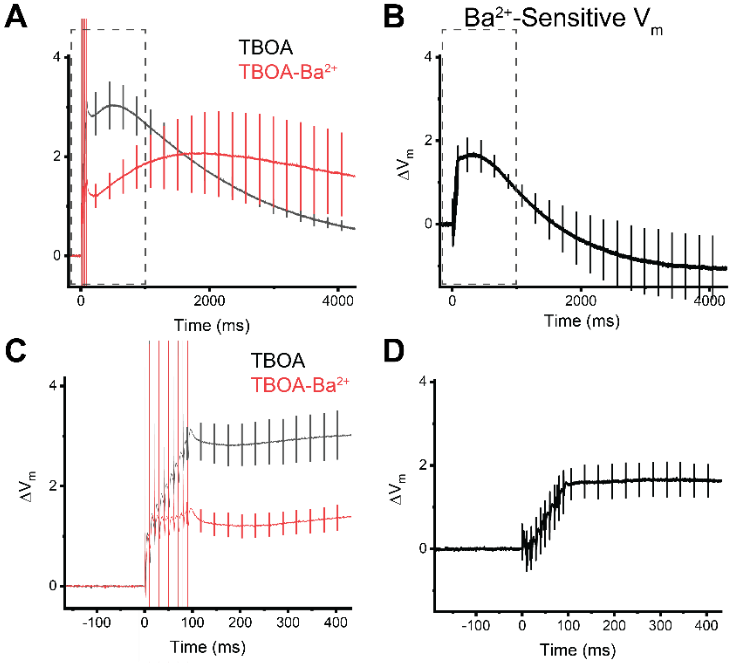


**Supplemental Figure 7: Kir4.1 depolarizes astrocyte soma during neuronal activity.** Astrocyte-whole cell current clamp recordings were made in the cortex to measure somatic $V_m$. In order to isolate the effects of Kir4.1 on astrocyte $V_m$ during neuronal activity, glutamate transporter activity was blocked with TFB-TBOA and responses to 10 stimuli at 100Hz were recorded before and after blockade of Kir4.1 with $Ba^{2+}$. A) Average paired traces before (black) and after (red) inhibition of Kir4.1 with $Ba^{2+}$, and B) the $Ba^{2+}$-sensitive $\Delta V_m$. These recordings show that Kir4.1 depolarizes astrocyte soma during neuronal activity. C and D) Expanded time scale (dashed boxes in A & B) to show $V_m$ during stimulus. N = 5 cells.

**Supplemental Figure 8**

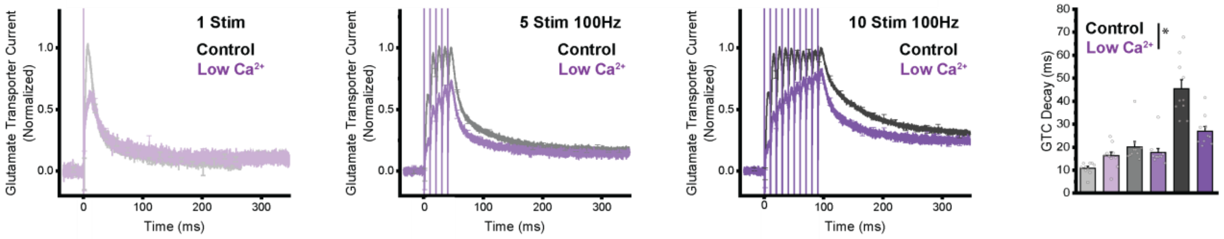


**Supplemental Figure 8: Low $Ca^{2+}$ Glutamate Transporter Currents.** Glutamate transporter currents were recorded from astrocytes with Control or Low $Ca^{2+}$ aCSF, showing enhanced glutamate clearance following trains of stimulation. Repeated measures ANOVA * = $p<0.05$. n = 10 cells/3 mice.

**Supplementary Methods:**

All animal protocols were approved by the Tufts Institutional Animal Care and Use Committee.

**Adeno-associated virus injection**

C57BL/6 or EAAT2-tdtomato(*10*) male and female mice (P30-45) were stereotaxtically injected with appropriate AAVs for the experiments in a single hemisphere with three injections sites (coordinates): (1.25, 1.25, 0.5), (1.25, 2.25, 0.5), and (1.25, 3.25, 0.5) (λ + *x*, +*y*, −*z*) mm. Mice were anesthetized with isoflurane for surgery, viruses were injected (1 µL per site (1:1 dilution with saline for single infection or 1:1 mix of viruses for co-infection, 0.15 µL/min) with $\sim 5 \times 10^9$ gene copies per virus. Mice were housed in 12/12 light/dark cycles following surgeries and were used for acute slice preparations 21–42 d following injection.

We would like to thank Dr. Loren Looger, Dr. Vincent Pieribone, Dr. Baljit Khakh, and Dr. Sergio Grinstein for making plasmids and constructs available that were used in this study.

| Virus | Source | AAV Production |
| --- | --- | --- |
| AAV5-GFAP-iGluSnFr(*24*) | Dr. Loren Looger | Addgene: 98930 |
| AAV5-hSyn-iGluSnFr(*24*) | Dr. Loren Looger | Addgene: 98929 |
| AAV5-GFAP-Archon1-EGFP (modified from Addgene 108417)(*7*) | Original plasmid from Dr. Edward Boyden | Vector Biolabs Custom Production |
| AAV5-GFAP-Arclight (modified from Addgene 36857)(*8*) | Original plasmid from Dr. Vincent Pieribone | Vector Biolabs Custom Production |
| AAV5-GFAP-Kir4.1-GFP(*18*) | Dr. Baljit Khakh | Duke Vector Core |
| AAV5-GFAP-Kir4.1-mCherry | Modified from Dr. Baljit Khakh's original construct | Duke Vector Core |
| AAV5-GFAP-tdtomato(*18*) | Dr. Baljit Khakh | Addgene 44332 |
| AAV5-GFAP-Lyn-pHluorin-mCherry (modified from Addgene 32002)(*13*) | Original plasmid from Dr. Sergio Grinstein | Vector Biolabs custom Production |

**Preparation of acute brain slices**

Cortical brain slices were prepared from AAV-infected mice(*4*). Mice were anesthetized with isoflurane, decapitated, and the brains were rapidly removed and placed in ice-cold slicing solution containing (in mM): 2.5 KCl, 1.25 $NaH_2PO_4$, 10 $MgSO_4$, 0.5 $CaCl_2$, 11 glucose, 234 sucrose, and 26 $NaHCO_3$ and equilibrated with 95% $O_2$:5% $CO_2$. The brain was glued to a Vibratome VT1200S (Leica Microsystems, Wetzlar, Germany), and slices (400 µm thick) were cut in a coronal orientation. Slices were then placed into a recovery chamber containing aCSF comprising (in mM): 126 NaCl, 2.5 KCl, 1.25 $NaH_2PO_4$, 1 $MgSO_4$, 2 $CaCl_2$, 10 glucose, and 26 $NaHCO_3$ (equilibrated with 95% $O_2$:5% $CO_2$). Slices were allowed to equilibrate in aCSF at 32°C for 1 hr. For astrocyte whole-cell experiments slices were loaded with sulforhodamine 101 (SR-101, 0.5 µm) in aCSF for 5 min at 32°C before equilibration(*36*) and were allowed to return to room temperature prior to electrophysiology/imaging. Low $Ca^{2+}$ aCSF reduced [$Ca^{2+}$] to 1mM and balanced divalents with increased [$Mg^{2+}$]. High [$K^+$] (7.5mM and 12.5mM) aCSF solutions were osmotically balance with substituting [$Na^+$].

### Live imaging

Arclight, Archon1, pHluorin, or iGluSnFr slices were placed into a submersion chamber (Siskiyou or Warner), held in place with small gold wires, and perfused with aCSF containing 20 μM DNQX (antagonist of AMPA receptors), and 50 μM APV (antagonist of NMDA receptors), equilibrated with 95% $O_2$:5% $CO_2$ and circulated at 2 ml/min at 34°C. A tungsten concentric bipolar stimulating electrode (FHC; Bowdoin, ME, USA) was placed in the deep cortical layers, and the upper cortical layers were imaged with a 60× water-immersion objective or 20X water-immersion objective (iglusnfr) (LUMPLANFL, Olympus) on a custom Prior Open-Scope with X-light V2 spinning disk confocal microscope (Crest-optics, 89North LDI). For two-stimulator experiments, a second identical tungsten stimulating electrode was placed in the upper cortical layers ~450 μm from the site of imaging. The 100 μs stimulus pulses were generated through a stimulus isolator ISO-Flex (A.M.P.I.). Stimulus intensity was set at 2X the resolvable threshold stimulation. Imaging was performed using a Prime95B (Photometrics) or Zyla5.5 (Andor) camera imaging, 12 bit digitization, 10 ms rolling shutter mode for 100 Hz temporal resolution, illuminated by a 470 or 640nm laser (89North) or LED (iGluSnFr, CoolLED) controlled by MicroManager(*37*). Arclight and Archon1 were imaged in confocal mode, using restricted illumination for enhanced laser illumination intensity, while iGluSnFr was imaged in bypass (epifluorescence) mode, using a GFP filter cube (Chroma). Stimulus responses were imaged in sequence of 5 repetitions of 1 Stim, followed by 5 and 10 at 100Hz Stim, followed by a no-stimulus control trace. This sequence was repeated >5 times for each slice. For each stimulus, a 1s burst was acquired (100 frames). For iGluSnFr imaging the same sequencing was used except that each stimulus condition was only repeated once in each sequence. Drugs were washed on for 5 min before recommencing imaging.

### Astrocyte whole-cell recordings

Astrocyte whole-cell recordings were performed similar to previous studies(*23, 38, 39*). Briefly, whole-cell patch-clamp recordings were made in an identical setup to the live-cell imaging. Astrocytes were identified by morphology (small, round cell bodies), membrane properties, and SR-101 labeling(*36*) as imaged with a Cy3 filter cube (excitation 560/40 nm, emission 630/75 nm, Chroma). Astrocyte internal solution contained the following (in mM): 120 potassium gluconate, 20 HEPES, 10 EGTA, 2 MgATP, and 0.2 NaGTP. The 4–12 MΩ borosilicate pipettes were used to establish whole-cell patch-clamp recordings using a Multiclamp 700B patch-clamp amplifier (Molecular Devices), sampled at 10 kHz using pClamp software. Once a whole-cell recording was established, cells were confirmed as astrocytes based on their passive membrane properties, low membrane resistance, and hyperpolarized resting membrane potential. Slices were perfused with aCSF containing 20 μM DNQX, and 50 μM APV, which was oxygenated and circulated at 2 ml/min at 34°C. Stimulus responses were activated as for live-cell imaging above. In voltage-clamp mode, whole-cell patch-clamped astrocytes were maintained at −80 mV. Changes in $V_m$ were recorded in current-clamp mode, while glutamate transporter currents were recorder in voltage-clamp mode. iGluSnFr decays were fit as previously with mono-exponential decay functions(*4*).

### NR2A-NMDA receptor EPSC current recording

NMDA receptor EPSCs were recorded similar to previous studies(*4, 39*). Briefly, whole-cell patch-clamp recordings were made in an identical setup to the live-cell imaging, with aCSF containing gabazine (10 μM), DNQX (20 μM), ifenprodil (5 μM), and D-serine (30 μM). Neurons identified by morphology in layer II/III of the cortex were whole-cell patch-clamped using 2–5 MΩ borosilicate glass electrodes containing the following (in mM): 120 D-gluconic acid, 120 CsOH, 10 HEPES, 10 EGTA, 0.5 $CaCl_2$, 20 TEA, 2 MgATP, and 0.2 NaGTP. In voltage-clamp mode, neurons were held at +40 mV to relieve the $Mg^{2+}$ block of the NMDA receptors and allowed to stabilize. NMDA currents were then evoked identically to the live cell imaging above. Access resistance was monitored throughout the experiment, and cells with more than a 25% change were excluded from analysis. NMDA current decays were fit as previously with mono-exponential decay functions(*4*).

**Immunofluorescence**

Fixed mouse brains were prepared by transcardial perfusion with 4% PFA. Fixed brains were sectioned at 30 µm using a Thermo Fisher Microm HM 525 cryostat. Brain sections were blocked using blocking buffer (5% normal goat serum, 1% BSA, in PBS) for 1 h at room temperature. NEUN (1:500, MAB377B, Millipore), GFP (1:500, ab13970 Abcam), glutamine synthase (1:500, MAB302, Millipore), Kir4.1 (1:200 APC-035 Alomone), GFAP (1:500 ab7260 Abcam) were diluted in PBS with 2% Triton X-100 and 5% blocking buffer. Cortical sections were incubated with diluted primary antibodies overnight at 4°C. Secondary antibodies (goat anti-rabbit Cy3, goat-anti chicken FITC, Jackson ImmunoResearch Laboratories) were diluted 1:500 in PBS with 5% blocking buffer and added to cortical sections for 2 h at room temperature. Slices were imaged with a Nikon A1R confocal microscope. Slices from 3 mice were stained for all experiments, with 2–4 slices per mouse visualized.

**Astrocyte territory segmentation**

Segmentation of the astrocytes labelled with tdTomato was performed in Imaris (Bitplane, Oxford Instruments) using the surface plug-in in a semi-automated manner. Astrocytes and their territory were carefully segmented using a watershed algorithm incorporated into the plug-in applied onto the tdTomato channel. The resulting segmented volume was used as a 3D mask to visualize and quantify the intensity of the Arclight channel.

**Imaging Analysis**

Analysis was performed using MATLAB (The MathWorks, Natick, MA) and Origin (Originlab, Northampton, MA). Following imaging, all acquired images from individual slices were aligned using the NoRMCorre algorithm(*40*). Following alignment, average $\Delta F/F_0$ images were generated and corrected for background fluorescence, using the no-stimulus and individual stimuli conditions. ROIs were generated using PCA/ICA algorithms(*14*), based on a spatial and temporal gaussian (1 pixel/frame kernel) filtered 10 Stimulation at 100Hz $\Delta F/F_0$ images. The analysis was given the full sweep duration and was blind to the stimulus timing. ROIs were spatially constricted to >2 pixels in size. The PCA/ICA identified the stimulus response from the full sweep and generated ROIs that were found throughout the image, had a non-random distribution, and had a similar kinetic profile to the $\Delta F/F_0$ signal seen across the entire image. To validate the approach, we compared the ROIs identified using PCA/ICA to an alternative approach in which pixels with the highest $\Delta F/F_0$ at the end of the stimulus were selected. The number of pixels selected was comparable to the number selected by PCA/ICA. This approach identified the noisiest pixels at the time of stimulation, had no kinetic similarity to the signal detected across the entire image, and were randomly distributed in space (Supp. Fig 7). Based on this comparison, PCA/ICA analysis was far superior. $\Delta F/F_0$ measures are susceptible to being distorted if the basal fluorescence ($F_0$) is very low. This concern is most relevant to the Archon GEVI which has a very dim baseline, while the Arclight GEVI is bright at baseline. To test whether the ROIs selected using PCA/ICA were biased toward low $F_0$ values, distorting $\Delta F/F_0$ measurements, we compared

ROI $\Delta F/F_0$ to $F_0$, on a hot spot-by-hot spot basis. This showed no correlation for either GEVI (Supp. Fig. 8). Lastly, the PCA/ICA algorithm finds similar responses for both Arclight and Archon1 (but in opposite directions) as Arclight is bright at baseline and reduces fluorescence, while Archon is dim at baseline and increase in brightness. Together this supports that the PCA/ICA-based approach to ROI detection is valid. Kymographs were generated by up-sampling ROIs 10x to enable sub-pixel alignment based on the center of mass of the detected ROI. Only ROIs less than 10 pixels area were included to avoid merged ROIs. For FWHM measurements individual slice kymographs were fits with a 2D gaussian function (Matlab) to calculate the FWHM. GEVI decays were quantified by calculating $T_{1/2}$ times based on the end of the stimulus.

### Pharmacology

Unless otherwise noted, all salts and glucose were obtained from Sigma-Aldrich. Drugs used in the study and their concentration: APV (NMDA antagonist, 50 μM, Tocris Bioscience)(*41*); DNQX (AMPA antagonist, 20 μM, Sigma)(*42*); D-Serine (NMDA coagonist, 30 μM, Sigma)(*43*); gabazine (6-imino-3-(4-methoxyphenyl)-1(6*H*)-pyridazinebutanoic acid hydrobromide, $GABA_A$ antagonist, 10 μM, Tocris Bioscience)(*44*); Ifenprodil ((1*S**,2*S**)-*threo*-2-(4-Benzylpiperidino)-1-(4-hydroxyphenyl)-1-propanol hemitartrate; NR2B antagonist, 5 μM, Tocris Bioscience)(*45*); TFB-TBOA (EAAT antagonist, 1 μM, Tocris Bioscience)(*46*); and TTX ($Na_V$ channel antagonist, 1 μM, Sigma). Drugs were kept as 1000× stock in $H_2O$ (APV, D-serine, gabazine, ifenprodil, TTX) or DMSO (DNQX, DPCPX, TFB-TBOA).
